# *OPRM1* A118G and serum β-endorphin interact with sex and digit ratio (2D:4D) to influence risk and course of alcohol dependence

**DOI:** 10.1101/200196

**Authors:** Jordan Bruno Gegenhuber, Christian Weinland, Johannes Kornhuber, Christiane Mühle, Bernd Lenz

## Abstract

Activation of mesolimbic mu-opioid receptor by its endogenous ligand, β-endorphin, mediates part of the rewarding effects of alcohol, yet there is controversial evidence surrounding the relationship between the functional mu-opioid receptor gene (*OPRM1*) A118G single nucleotide polymorphism and alcohol dependence risk. Some preclinical evidence suggests that sex and sex hormone-dependent prenatal brain organization may interact with the opioid system to influence alcohol drinking behavior. We genotyped 200 alcohol-dependent patients and 240 healthy individuals for the A118G variant and measured serum β-endorphin level at recruitment and during acute withdrawal. We then evaluated the association between these factors and alcohol dependence risk and outcome in the context of both sex and second-to-fourth digit length ratio (2D:4D) – a biomarker of prenatal sex hormone load. For the first time, the AA genotype was found to be associated with elevated alcohol-related hospital readmission risk, more readmissions, and fewer days until first readmission in male but not female patients. Upon accounting for 2D:4D, the G-allele predicted alcohol dependence and more readmissions (1 vs ≥2) in males, suggesting prenatal sex hormones interact with *OPRM1* to influence addiction pathology. Withdrawal β-endorphin level also correlated negatively with withdrawal severity in females but not in males, indicating β-endorphin might protect against withdrawal-induced stress in a sex-specific manner. Organizational effects of sex hormones may prime individuals for alcohol dependence by inducing permanent changes to the endogenous opioid system.

## 1. Introduction

Activation of mesolimbic mu-opioid receptor (MOR) by its primary endogenous ligand, β-endorphin (β-END), has been shown to mediate part of the rewarding effects of alcohol via stimulating nucleus accumbens dopamine release (Heilig et al., 2011). For this reason, MOR and β-END are considered high-interest pharmacological targets for the treatment and prevention of alcohol dependence and relapse. The relatively common single nucleotide polymorphism (SNP) rs1799971 within exon 1 of the mu-opioid receptor gene (*OPRM1*), in which an adenine-guanine transition (A118G) encodes for an asparagine-aspartic acid substitution (N40D), has been studied in the context of alcohol dependence due to its strong yet opposing effects on receptor function and expression. The G-allele has been reported to increase binding affinity for β-END three-fold (Bond et al., 1998) and cause a four-fold increase in ventral striatal dopamine release following alcohol stimulation (Ramchandani et al., 2011) but also reduce *OPRM1* mRNA and protein level in human post-mortem brain tissue and transfected Chinese hamster ovary cells (Zhang et al., 2005).

Despite these potent functional effects, previous meta-analyses have been unable to establish a clear association between *OPRM1* A118G and alcohol dependence in humans (Chen et al., 2012; Schwantes-An et al., 2016). Several studies have found the G-allele to be associated with elevated dorsal striatal cue-reactivity (Bach et al., 2015), craving (van den Wildenberg et al., 2007), and subjective feelings of stimulation and happiness (Ray et al., 2004; 2013) following alcohol consumption. However, epidemiological research and meta-analyses have linked both the A- and G-allele to increased risk of alcohol dependence amongst European cohorts consisting of both sexes (Bart et al., 2005; Schwantes-An et al., 2016). This inconsistency may be due to an inability or failure to account for certain confounding factors.

There is evidence suggesting that sex-*OPRM1*, sex-β-END, and prenatal sex hormone-*OPRM1* interactions may influence addiction pathology. Barr et al. (2007) found that the G-allele of the primate equivalent of the A118G SNP significantly increases alcohol-induced stimulation, ethanol consumption, alcohol preference, and percentage of days intoxicated in male but not female rhesus macaques. In a humanized mouse model carrying this polymorphism, male GG mice self-administered more nicotine than male AA mice, whereas there was no difference in female animals (Bernardi et al., Gegenhuber et al. *OPRM1*, β-endorphin, and alcohol dependence 4 2016). A different mouse model containing a point mutation (A112G) equivalent to the human A118G SNP also revealed male GG mice tend to have a higher conditioned place preference for a morphine-paired environment than male AA mice (Mague et al., 2009). In humans, naltrexone, a MOR antagonist, significantly and dose-dependently reduces the number of heavy drinking days in males but not in females (Garbutt et al., 2005). However, few studies have investigated sex-specific *OPRM1* genotype effects on alcohol dependence in a clinical cohort. Based on preliminary evidence, we hypothesize that the A118G SNP as well as three additional *OPRM1* promoter variants relate more strongly to alcohol dependence risk and outcome in males or females than in the combined cohort. Furthermore, since male heavy drinker plasma β-END level has been shown to be higher than in female heavy drinkers (Gianoulakis et al., 2003), peripheral β-END levels in patients, before and during withdrawal, and in healthy individuals were investigated on a sex-specific basis.

Recent evidence has also revealed an interaction between prenatal testosterone level and *OPRM1* influencing alcohol consumption in rodents (Huber et al., in press). In humans, elevated exposure to androgens during the prenatal development window has been associated with increased risk of alcohol dependence (Kornhuber et al., 2011; Lenz et al., 2017) and other behavioral disorders such as video game addiction (Kornhuber et al., 2013). From a mechanistic standpoint, sex hormones have been shown to induce permanent, organizational changes to the brain’s reward system at the neuronal and molecular level (Brown et al., 2015). Huber et al. (in press) found in mice that prenatal treatment with flutamide, a potent androgen receptor antagonist, caused a significant reduction in adult ventral striatal *OPRM1* RNA and a significant increase in alcohol self-administration. Given this finding, we decided to also investigate prenatal sex hormone-*OPRM1* effects on alcohol dependence and outcome, using the second-to-fourth digit length ratio (2D:4D) as a biomarker for prenatal sex hormone load (Berenbaum et al., 2009; Zheng and Cohn, 2011).

## 2. Methods

### 2.1 Cohort characteristics

The individuals included in this investigation were recruited between 2013 and 2014 for the **N**eurobiology **o**f **A**lco**h**olism (NOAH) study at the Universitätsklinikum Erlangen Department of Psychiatry and Psychotherapy and the Klinikum am Europakanal Clinic for Psychiatry, Psychotherapy, and Psychosomatic Medicine in Erlangen, Germany (Lenz et al., 2017). 200 early-abstinent, alcohol-dependent patients (113 males and 87 females) admitted as inpatients for withdrawal treatment were selected through an intensive screening process, and 240 healthy controls (133 males and 107 females) were selected after two telephone screenings and an on-site interview, in which individuals with prior psychiatric inpatient treatment and/or any psychiatric outpatient treatment during the past ten years were excluded. Median age (years) for both patients and controls was 48 (interquartile ranges [IQR] 42/54 and 39/56) (Mann-Whitney *U* test [MWT], p > 0.05), and median body mass index (kg/m^2^) was 24.7 (IQR 22.1/28.2) and 26.5 (IQR 23.5/29.3) (MWT, p > 0.05), respectively. In the patient group, median lifetime drinking (kg) and daily ethanol intake (g/d since onset) were 483 (IQR 270/1,195) and 120 (IQR 58/240). Whole blood, behavioral scores, and other parameters (Lenz et al., 2017) were collected at time of recruitment (day A) in patients and controls and a median of five days later (day B) in patients only, during which they underwent withdrawal. The German version of the Clinical Institute Withdrawal Assessment for Alcohol revised (CIWA-Ar) scale was used to measure alcohol withdrawal severity (Stuppaeck et al., 1994). Blood samples were collected in the morning for all individuals to minimize circadian effects on hormone level. Individuals' hands were scanned using the HP Scanjet G4050 (Hewlett-Packard, Palo Alto, CA, USA), and 2D:4D was calculated by three blinded raters, who measured the absolute lengths of the second and fourth digits (GNU Image Manipulation Program, www.gimp.org). Lower 2D:4D indicates higher prenatal androgen and lower estrogen load, and higher 2D:4D suggests the opposite. Medical records for each patient were accessed 24-months after study recruitment to investigate the number of alcohol-related hospital readmissions and days until first readmission.

### 2.2 Blood preparation

After collection, blood samples were centrifuged (10 min, 2000 g at room temperature), and serum was placed into storage at -80°C. Genomic DNA (gDNA) was extracted from whole blood using the Gentra Puregene Blood Kit (Qiagen, Venlo, Netherlands) and stored at 4°C.

### 2.3 Genotyping

Genotyping for three *OPRM1* SNPs (rs1799971 [A118G], rs3798677, rs3798678) was performed by high resolution melting (HRM) of polymerase chain reaction (PCR) products in the Roche LightCycler 480 II (Roche Holding AG, Basel, Switzerland). The PCR primer sequences are listed in Table 1. The reaction mixture consisted of 1x Rovalab buffer and 0.1 U Taq polymerase (Rovalab GmbH, Teltow, Germany), 0.03 μl CyGreen (1:100 dilution, Enzo Life Sciences Inc., Farmingdale, NY, USA), 10 ng gDNA, 1.5-3.0 mM MgCl2 (Table 1), 200 μM dNTPs (each), and 200 nM forward and reverse primers (Sigma-Aldrich, St. Louis, MO, USA) in a total volume of 10 μl. PCR conditions were: 2 min at 95°C followed by 40 x (15s at 95°C, 30s at primer-specific annealing temperatures [Tab. 1] 15s at 72°C). After PCR, product denaturation was performed at 95°C for one minute, followed by rapid cooling to 40°C. A melting curve was generated by heating at 0.02°C/s from 70°C to 90°C, and analysis was performed using the Roche LightCycler 480 Gene Scanning Software v1.5. Genotypes for the three SNPs were confirmed by Sanger sequencing of gDNA samples selected to act as standards (AA, AG, GG for each SNP).

**Table 1.**
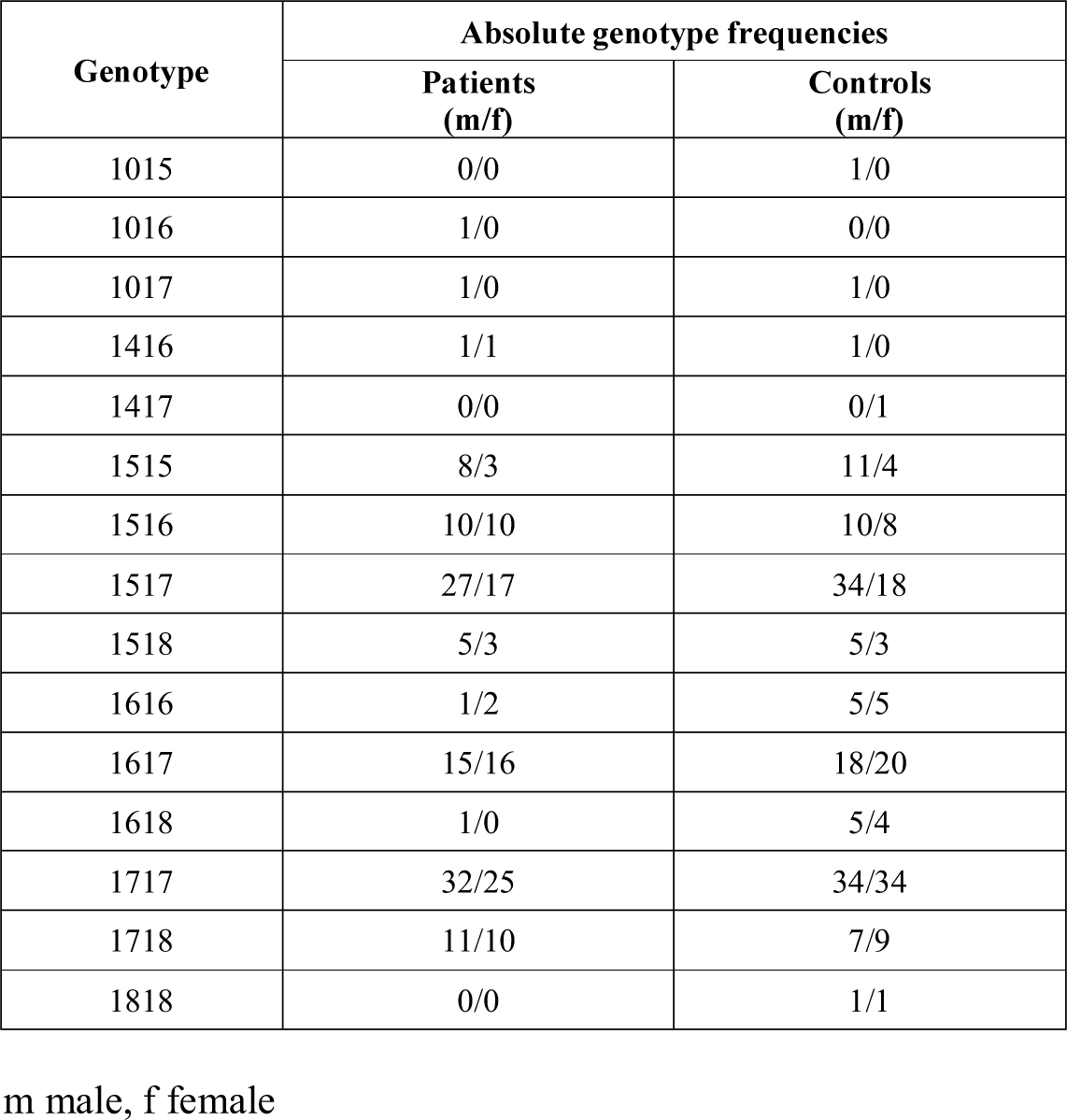
RefSNP numbers and PCR conditions for the three SNPs genotyped in this study. rs3798677 and rs3798678 are silent variants located within the *OPRM1* promoter.

**Table 2.**
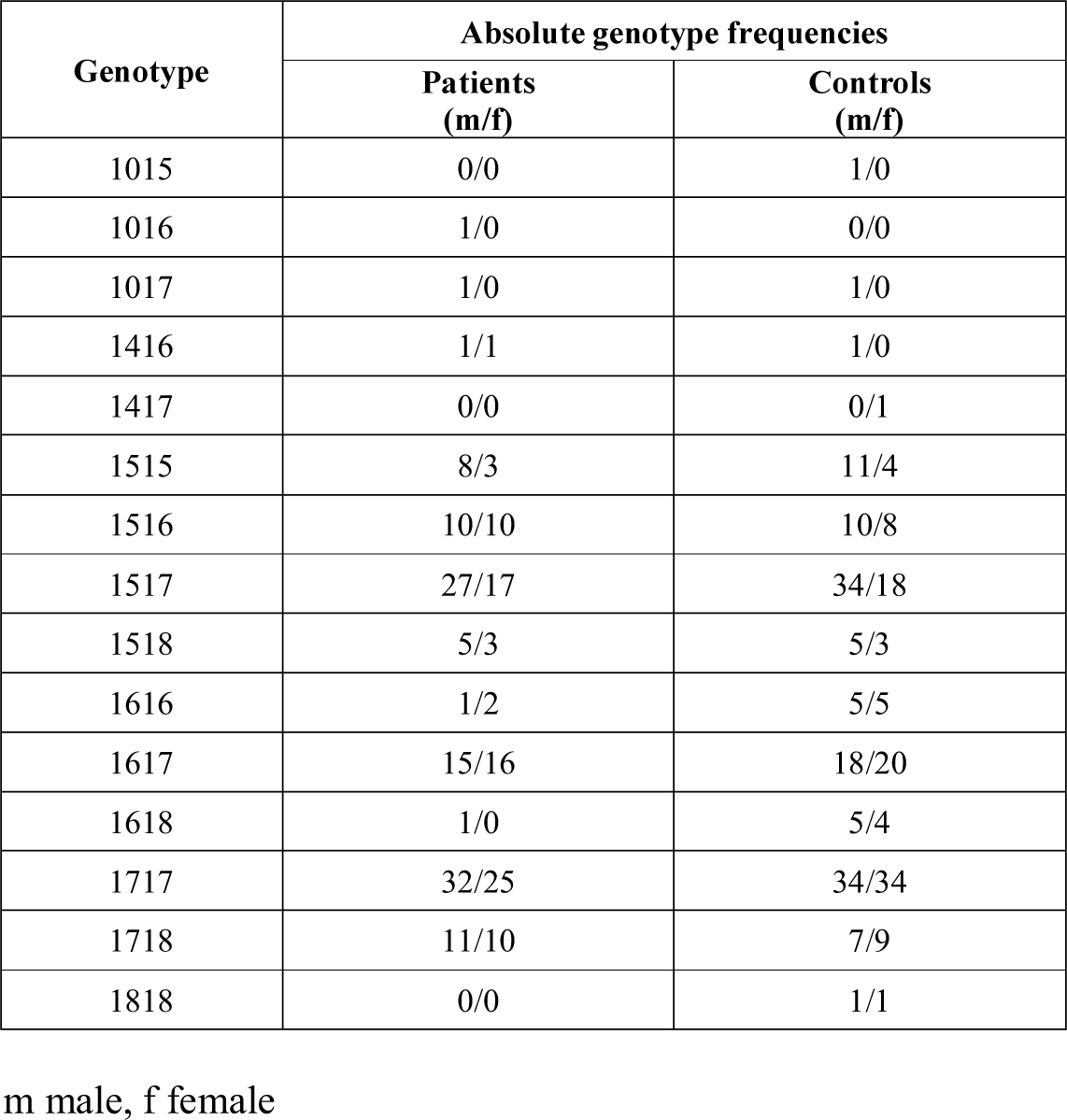
Absolute *OPRM1* CAn genotype frequencies.

Genotyping for the *OPRM1* CAn dinucleotide repeat polymorphism, described in Kranzler et al. (1998), was performed by fragment length analysis of PCR products in an Applied Biosystems 48-capillary array (Thermo Fisher Scientific, Waltham, MD, USA). The PCR primer sequences were the same as for genotyping rs3798678 except the forward primer was fluorescently tagged with 6-fluorescein amidite (Sigma-Aldrich, St. Louis, MO, USA). PCR reaction mixture and conditions were the same as for the rs3798678 HRM-PCRs, except for the absence of the CyGreen intercalating dye. Following PCR, products were diluted 1:20 in MilliQ water, and 1 µl of the dilution was mixed with 10 µl formamide (Sigma-Aldrich) and 0.5 ul GeneScan 500 ROX dye size standard (Thermo Fisher Scientific) prior to fragment length analysis. CAn genotypes were confirmed by Sanger sequencing.

10% of all samples' genotypes for the polymorphisms were replicated with 100% accordance. The four genetic variants were in Hardy-Weinberg equilibrium within the NOAH cohort (rs1799971 [A118G]: χ^2^ = 0.180, p = 0.672; rs3798677: χ^2^ = 1.521, p = 0.217; rs3798678: χ^2^ = 1.521, p = 0.217; CAn: χ^2^ = 13.550, p = 0.809).

### 2.4 β -END quantification by enzyme-linked immunosorbent assay (ELISA)

Serum β-END was quantified using the Endorphin, beta (Human) EIA Kit by Phoenix Pharmaceuticals, Inc. (Burlingame, CA, USA) following the manufacturer's guidelines. 25 µl of original, unpurified serum were assayed in duplicate, and peptide concentrations were extrapolated from a standard curve (7 dilutions from 30 ng/ml to 0.01 ng/ml), which was run on every 96-well plate to minimize risk of interplate variation. The intra-assay and inter-assay coefficients of variation were 7% and 14%, respectively.

### 2.5 Statistics

Alpha (two-tailed) was set to 0.5. If not otherwise stated, we report median and IQR. For group comparisons, we used MWTs, Kruskal-Wallis tests (KWTs), and Wilcoxon signed-rank tests. The Spearman's method was employed to evaluate bivariate correlations and χ^2^ tests with odds ratios (OR) for differences in the frequencies. Log-rank [Mantel-Cox] tests were used for survival curves. To increase statistical power, heterozygous subjects were combined with homozygous minor allele individuals for SNP analysis (AA study subjects vs. G-allele carriers). In case of missing data points, study subjects were excluded from the specific analyses, and the exact case number is stated. Data were analyzed using IBM SPSS Statistics Version 21 for Windows (SPSS Inc., Chicago, IL, USA) and Graph Pad Prism 5 (Graph Pad Software Inc., San Diego, CA, USA).

## 3. Results

### 3.1 Trend for an association between OPRM1 A118G and alcohol dependence risk in females

The *OPRM1* A118G genotype (AA vs. AG/GG) did not predict risk of alcohol dependence in the total cohort (χ^2^test; OR = 1.12 [95%-confidence interval (95%-CI) 0.71-1.76], χ^2^ = 0.233, p = 0.629) or the male subgroup (OR = 0.81 [95%-CI 0.45-1.44], χ^2^ = 0.528, p = 0.467). However, there is a trend for AA females to have an elevated risk (OR = 1.91 [95%-CI 0.89-4.06], χ^2^ = 2.848, p = 0.091).

### 3.2 Sex-specific association between OPRM1 A118G and prospective alcohol-related hospital readmission

*OPRM1* A118G significantly predicted the outcome of male patients but not of female patients over the 24-month period following study recruitment. In males, the AA genotype was significantly associated with an elevated risk of alcohol-related hospital readmission (Males, readmitted: n(AA) = 62, n(AG/GG) = 14, non-readmitted: n(AA) = 21, n(AG/GG) = 16; females, readmitted: n(AA) = 38, n(AG/GG) = 8, non-readmitted: n(AA) = 37, n(AG/GG) = 4; Fig. 1A), higher median number of readmissions (Fig. 1B), and fewer mean days until first readmission (Fig. 1C). No significant associations between genotype and outcome were observed in females (readmitted: n(AA) = 38, n(AG/GG) = 8, non-readmitted: n(AA) = 37, n(AG/GG) = 4; Fig. 1A-B, D).

**Figure 1.**
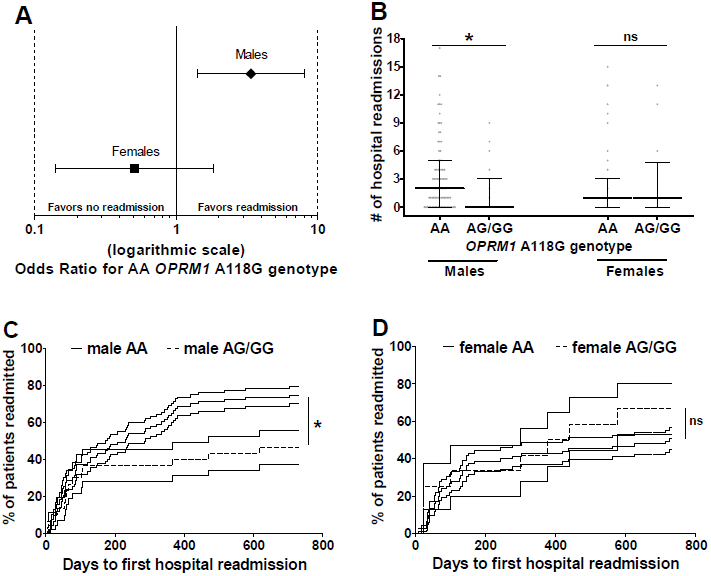
Effect of ***OPRM1*** A118G genotype on prospective alcohol-related hospital readmission during the 24-month follow-up. (A) A118G AA genotype is associated with elevated readmission risk in males (OR = 3.37 [95%-CI 1.41-8.07], χ^2^ = 7.863, p = 0.005) but not in females (OR = 0.51 [95%-CI 0.14-1.85], χ^2^ = 1.063, p =0.303). (B) AA males had a higher median number of readmissions than male G-allele carriers (AA: 2 [IQR 0-5], AG/GG: 0 [IQR 0-3], Mann-Whitney U test, *U* = 869.0, p = 0.012), whereas genotype did not predict number of readmissions in females (AA: 1 [IQR 0-3], AG/GG: 1 [0-5], MWT, *U* = 399.5, p = 0.509). Data are presented as median ± IQR. (C) AA males had fewer mean days until first readmission than male G-allele carriers (AA: 313.3 ± 30.6, AG/GG: 457.7 ± 57.9, Log-rank [Mantel-Cox] test, χ^2^ = 5.562, p = 0.018). (D) There was no difference in days until first readmission between AA females and female G-allele carriers (AA: 447.7 ± 35.9, AG/GG: 398.8 ± 83.0, Log-rank [Mantel-Cox] test, χ^2^ = 0.913, p = 0.339). C and D show survival curves ± SEM. OR odds ratio, 95%-CI 95%-confidence interval. *p < 0.05, ns not significant.

### 3.3 No significant associations of rs3798677, rs3798678, and CAn polymorphisms with alcohol dependence or outcome

10 and 14-18 CAn repeats were detected within the NOAH cohort (Tab. 2). CAn allele length and genotype were investigated for associations with alcohol dependence or outcome in the NOAH cohort. Mean CAn allele length did not significantly predict risk of alcohol dependence or alcohol-related hospital readmission (MWTs), nor was it significantly correlated with number of readmissions or days until first readmission (Spearman-Rho) in both the total group and the male and female subgroups. Similarly, no significant associations were observed between CAn genotype and alcohol dependence or outcome (χ^2^ tests, KWTs).

Promoter region SNPs rs3798677 (-73 bp relative to CAn) and rs3798678 (-3 bp relative to CAn) had identical genotypes in the cohort (Tab. 1) and were also found to be in strong linkage disequilibrium with A118G (rs3798677 AA/G-allele carriers vs. A118G AA/G-allele carriers; χ^2^ = 8.287, p = 0.004). Similar to the CAn promoter variant, rs3798677 and rs3798678 did not significantly predict dependence or outcome and were also not associated with the number of readmissions or days until first readmission (χ^2^ tests, MWTs, Log-rank [Mantel-Cox] tests).

### 3.4 Serum β-END level declines during acute withdrawal in male and female patients and relates to withdrawal severity

At baseline, there was no significant difference in median serum β-END level between patients and controls in both males and females. During a median of five days withdrawal, patient β-END concentrations declined significantly to a level that was also significantly lower than in control subjects (Fig. 2A). Moreover, there was a significant negative correlation between withdrawal serum β-END level (day B) and CIWA-Ar withdrawal severity score in female patients (Fig. 2B) but not in males (Spearman-Rho, n = 82, ρ = -0.097, p = 0.385). No significant associations between patient serum β-END level and outcome were detected, except for one finding that day B β-END level was correlated with days until first readmission in males (Spearman-Rho, n = 94, ρ = -0.232, p = 0.025).

**Figure 2.**
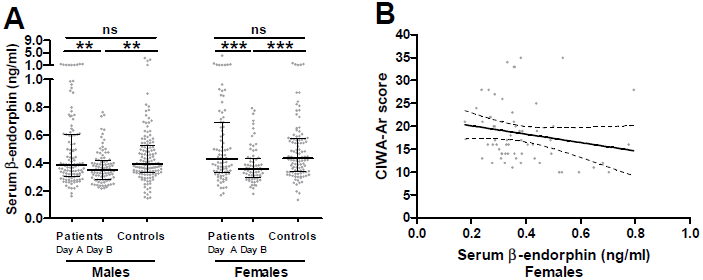
Sex-specific serum β -END level in study subjects and correlation of day B β -END level with withdrawal severity in female patients. (A) There was no difference in serum β-END level between patients and controls at baseline (male patients: 0.388 ng/ml [IQR 0.307-0.607], male controls: 0.392 ng/ml [IQR 0.330-0.526], Mann-Whitney *U* test, *U* = 7507.0, p = 0.989; female patients: 0.428 ng/ml [IQR 0.331-0.692], female controls: 0.433 ng/ml [IQR 0.335-0.574], Mann-Whitney *U* test, *U* = 4638.0, p = 0.966). During withdrawal of a median of 5 days, patient serum β-END level declined (Wilcoxon signed-rank tests, male patients: z = -3.2, p = 0.001, female patients: z = -3.5, p < 0.001) to a lower level than in control subjects (male patients: 0.348 ng/ml [IQR 0.280-0.415], Mann-Whitney *U* test, *U* = 4627.0, p = 0.001; female patients: 0.356 ng/ml [IQR 0.297-0.435], Mann-Whitney *U* test, *U* = 2525.0, p < 0.001). Data are shown as median ± IQR. Two outliers – one male and one female control – above 10 ng/ml β-END level are not shown. (B) Serum β-END level on day B correlated significantly with severity of alcohol withdrawal syndrome, quantified by the CIWA-Ar score (Spearman-Rho, n = 62, ρ = -0.282, p = 0.026). Dotted lines represent the 95% confidence intervals of the best-fit from a linear regression analysis. CIWA-Ar Clinical Institute Withdrawal Assessment for Alcohol revised scale, IQR interquartile range, **p = 0.001, ***p < 0.001, ns not significant.

### 3.5 A118G-2D:4D interaction influences risk and outcome of alcohol dependence

To test whether AA study subjects differ from G-allele carriers with regard to the effect of prenatal sex hormone load on risk of alcohol dependence, we sex- and genotype-specifically normalized patients' 2D:4D values to the mean 2D:4D and its standard deviation of the control group (= [each patient's 2D:4D value – mean controls' 2D:4D value] / standard deviation of controls' 2D:4D values). Male G-allele carrier patients differed significantly from AA male patients with regard to the normalized 2D:4D values (Fig. 3A), meaning a significantly stronger 2D:4D deviation between patients and controls in the G-allele carrier group than in the AA group. No significant 2D:4D-A118G interaction was observed in females (Fig. 3B). Subsequently, we explored whether similar interactions are also relevant to the patients' outcome. We previously reported a significant negative correlation between 2D:4D values and number of alcohol-related hospital readmissions during the 12-month follow-up in individuals with at least one readmission (Lenz et al., 2017). Thus, we sex- and genotype specifically normalized 2D:4D values of each patient with at least two readmissions to the mean 2D:4D and standard deviation of patients with only one alcohol-related readmission during the follow-up (= [2D:4D value of each patient with one readmission – mean 2D:4D values of patients with at least two readmissions] / standard deviation of 2D:4D values of patients with at least two readmissions). In support of the previous finding, male G-allele carrier patients deviated significantly from AA male patients with regard to the normalized 2D:4D values (Fig. 3C), meaning a significantly stronger 2D:4D deviation between patients with one readmission and patients with at least two readmissions in the G-allele carrier group than in the AA group. No significant 2D:4D-A118G interaction with respect to alcohol-related readmission was found in females (Fig. 3D). These two observations suggest that in males, higher prenatal androgen (and lower estrogen) load interacts with the *OPRM1* A118G G-allele to elevate risk and worsen outcome of alcohol dependence.

**Figure 3.**
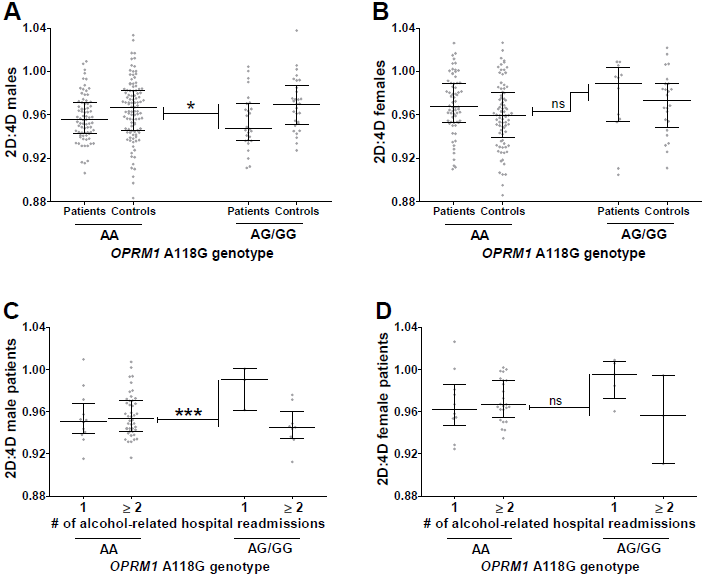
Sex-specific interaction of *OPRM1* A118G genotype with 2D:4D in risk and course of alcohol dependence. (A) We sex- und genotype-specifically normalized patients' 2D:4D values to the mean 2D:4D and standard deviation of the control group and found that male G-allele carrier patients deviated significantly from AA males with regard to the normalized 2D:4D values (normalized 2D:4D; AG/GG: n = 27, median = -0.941, AA: n = 76, median = -0.331, Mann-Whitney *U* test, *U* = 742.0, p = 0.033). (B) This difference was not observed in females (normalized 2D:4D; AG/GG: n = 12, median = 0.639, AA: n = 67, median = 0.240, Mann-Whitney *U* test, *U* = 375.0, p = 0.712). (C) 2D:4D value of each patient with at least two readmissions was sex- and genotype-specifically normalized to the mean 2D:4D and standard deviation of patients with one readmission; these normalized values differed significantly between G-allele carriers and AA study subjects in males (normalized 2D:4D; AG/GG: n = 9, median = -1.922, AA: n = 44, median = -0.020, Mann-Whitney *U* test, *U* = 23.0, p <0.001) but not in females (D; normalized 2D:4D; AG/GG: n = 3, median = -1.771, AA: n = 24, median = -0.025, Mann-Whitney *U* test, *U* = 14.0, p = 0.090). Data shown as median ± interquartile range. *p < 0.05, *** p < 0.001, ns not significant.

## 4. Discussion

Over the last 30 years, the proportion of neuroscience studies including both sexes has grown substantially, yet approximately 80% of such publications still fail to analyze results by sex despite clear sex differences in the risk and pathology of most major neuropsychiatric disorders, including alcohol dependence (Beery and Zucker, 2011). The present study aimed to investigate sex-specific *OPRM1* and β-END effects on alcohol dependence risk and outcome as well as prenatal sex hormone-*OPRM1* interactions.

In the NOAH cohort, the *OPRM1* A118G SNP did not predict alcohol dependence in either males or females but related sex-specifically to outcome. To our knowledge, this is the first publication to demonstrate that the AA genotype is associated with increased risk of alcohol-related hospital readmission, more readmissions, and fewer days until first readmission in male patients and therefore may be relevant for predicting relapse in a clinical setting. In previous naltrexone studies, Oslin et al. (2003) and O'Malley et al. (2008) found that placebo-treated AA alcohol-dependent patients were more likely to relapse than G-allele carriers, but this association did not achieve statistical significance – perhaps because a large proportion of the two study cohorts consisted of females. Schacht et al. (2017) also did not observe a significant difference in percentage of heavy drinking days between AA individuals and G carriers, although the duration of their study (10 months) was shorter than ours (24 months). Considering the G allele has been shown to strongly increase MOR binding affinity for β-END, we hypothesize the AA genotype elevates readmission risk by hyposensitizing MOR, causing dependent patients to consume more alcohol in order to overcome intrinsic reward deficit (Kleinjan et al., 2015).

Interestingly, we found that the A118G polymorphism does predict alcohol dependence in male patients upon accounting for 2D:4D, suggesting *OPRM1* and sex hormones interact in the prenatal state to influence addiction risk in adult life. Additionally, 2D:4D and *OPRM1* A118G interact to predict more readmissions (1 vs ≥2) in male patients; the group of patients with no recorded alcohol-related readmissions was excluded because it does not account for possible hospital readmissions outside the Erlangen area. The hypothesized interaction between prenatal sex hormone load and MOR signaling has recently been confirmed in a rodent study, in which intraperitoneal injection of pregnant dams with the potent androgen receptor antagonist flutamide resulted in significantly reduced ventral striatal *OPRM1* RNA expression and reduced alcohol consumption in the adult male offspring (Huber et al., in press). Other studies indirectly corroborate this finding. Prenatal stress, which is associated with children's 2D:4D, has been shown to influence adult pain sensitivity in a sex-specific manner (Butkevich et al., 2007), and prenatal exposure to cigarette smoke – another contributor to prenatal androgen load (Lenz et al., 2017) – reduces adult ventral striatal reward response (Müller et al., 2013) and also interacts with the *OPRM1* rs2281617 SNP to elevate adolescent dietary fat preference (Lee et al., 2015). Our findings in combination with these studies suggest organizational sex hormone signaling may induce lasting changes to the endogenous opioid system and mesolimbic reward pathway.

The functional effects of the A118G polymorphism may explain how 2D:4D and *OPRM1* A118G interact to influence alcohol dependence. Previously, the G-allele was shown to confer both activational and inhibitory effects on mesolimbic reward signaling in that it increases receptor binding affinity for β-END but also impairs *OPRM1* transcription, making it an unreliable predictor of alcohol dependence. However, after integrating 2D:4D values in our analyses, the G-allele becomes significantly associated with elevated risk, as prenatal androgens have been shown to increase *OPRM1* RNA expression irrespective of genotype and thus eliminate the protective, inhibitory effects of the G-allele on receptor expression (Huber et al., in press). This 2D:4D-*OPRM1* A118G interaction may also explain why the G-allele tends to protect against dependence risk in our cohort of female study subjects (OR = 1.91, p = 0.091) – females are exposed to far lower amounts of prenatal sex hormones than males and therefore should retain the protective, inhibitory effects of the G-allele. Additional *in vitro* and animal experiments will be required to establish the exact mechanism of the prenatal sex hormone-*OPRM1* interaction, although androgen- and estrogen-induced epigenetic modifications in the form of DNA methylation may be a promising starting point (Ghahramani et al., 2014).

In addition to *OPRM1* variation, we measured serum β-END level in healthy subjects and alcohol-dependent patients at baseline and during early alcohol withdrawal. At time of recruitment, patients' serum β-END did not significantly differ from controls' but strongly declined during withdrawal in both males and females, which is consistent with previous studies in males (Esel et al., 2001; Marchesi et al., 1996) and reveals this effect to be unrelated to sex. This finding indicates that both male and female patients may depend on alcohol to maintain a homeostatic serum β-END level.

At this time, it is unclear how β-END relates to withdrawal. In the NOAH cohort, β-END level was not associated with follow-up outcome in either sex, except for one questionable finding that may be a false positive. The lack of association may be due to several reasons. Previous studies have found peripheral β-END and brain β-END to be under independent regulation, so our serum measurements may not accurately represent the effects of withdrawal on brain β-END processing (Veening et al., 2012). Also, in our analyses, we have compared β-END level during acute withdrawal with long-term outcome when it is unclear whether β-END is stable or dynamic throughout the withdrawal period. β-END level after longer withdrawal intervals may prove to be a stronger predictor of long-term outcome.

Interestingly, in a study completed by Racz et al. (2008), female mice tended to be more affected by β-END deficiency with regard to physical withdrawal severity than male mice – an observation which validates our finding that CIWA-Ar withdrawal severity score correlates negatively with serum β-END concentration during withdrawal in females but not in males. Both β-END deficiency and female sex contribute to a higher functioning HPA axis (Goel et al., 2014). Thus, endogenous opioid production and biological sex may converge on the HPA axis to influence withdrawal-induced stress – an interaction that further highlights sex differences in the etiology of alcohol dependence and outcome. Sex hormone signaling may also exert direct effects on β-END production. A previous study from our group found that the number of CAGn repeats within exon 1 of the androgen receptor (AR) gene correlates negatively with promoter methylation of the β-END precursor gene – pro-opiomelanocortin – and that this methylation is partly responsible for the association between the AR CAGn variant and alcohol craving (Muschler et al., 2014). Understanding these sex effects is of high clinical interest, as it may shed light on the mechanisms of current withdrawal treatments, such as acamprosate, which has been found to both improve withdrawal symptoms and reverse withdrawal β-END deficiency (Hammarberg et al., 2009; Zalewska-Kaszubska and Czarnecka, 2005).

There are several limitations to our approach. The associational study design prohibits us from establishing a causal role for prenatal sex hormones in influencing *OPRM1*. Our assessment of 2D:4Dgenotype interactions in predicting outcome is limited by the low number of G-carriers who readmitted to the hospital. We also have not corrected for multiple hypothesis testing. Furthermore, prenatal sex hormone load is measured indirectly using 2D:4D as a biological proxy, although it is quite reliable (Zheng and Cohn, 2011).

Previous studies report controversial results regarding the role of the *OPRM1* A118G SNP in alcohol dependence. Here we show that upon analyzing findings by sex, AA genotype significantly predicts alcohol-related hospital readmission, number of readmissions, and days until first readmissions in male patients. Additionally, upon accounting for sex and 2D:4D – a biomarker for prenatal sex hormone load – the *OPRM1* G-allele significantly predicts alcohol dependence and outcome. Serum β-END level during withdrawal was also found to relate sex-specifically to outcome, as it correlates negatively with CIWA-Ar withdrawal severity in females but not in males. Since this study is limited by its associational design, future work is needed to assess whether prenatal sex hormones influence alcohol dependence and outcome by directly influencing *OPRM1* and β-END or whether prenatal sex hormones and the opioid system are independent predictive factors of risk. However, given the existing wealth of information on the cross-regulation between opioid and sex hormone signaling, the observations collected here support a novel mechanism of addiction pathogenesis.

## Declarations

### Ethics approval and consent to participate

The NOAH study was previously approved by the Ethics Committee of the Medical Faculty of the Friedrich-Alexander University Erlangen-Nürnberg (NOAH study ID 81_12) and was in accordance with the sixth revision of the Declaration of Helsinki, set forth by the World Medical Association (Seoul, 2008), as well as the International Conference on Harmonization Guidelines for Good Clinical Practice (1996).

## Conflict of Interest

The authors do not declare any personal or financial conflicts of interest.

## Funding

This work was supported by intramural grants from the University Hospital of the Friedrich-Alexander University Erlangen-Nürnberg (FAU) and the Fulbright U.S. Student Program. The funders had no role in the study design, data collection, analysis, decision to publish, or preparation of the manuscript.

## Authors' contributions

JBG, CM, and BL designed the study. JBG performed the *OPRM1* variant genotyping, and CM and BL performed the serum β-END ELISAs. CW and BL checked patients' medical records for alcohol-related hospital readmission. JBG and BL performed the statistical analyses and wrote the first draft of the paper. All authors contributed in a substantial way to the project, provided feedback on the manuscript, and approved the final version for publication.

## Acknowledgment

We thank the German-American Fulbright Kommission and the Institute of International Education for selecting this project for funding. We gratefully appreciate the support of Dr. Birgit Braun, Juliane Behrens, Franziska Kreß, Sarah Kubis, Katrin Mikolaiczik, Juliana Monti, Marcel-René Muschler, Hedya Riesop, Sarah Saigali, Marina Sibach, and Petya Tanovska in recruiting patients, technical support, and 2D:4D quantification. We also thank Dr. Andreas Ahnert, Ute Hamers, and Dr. Kristina Bayerlein for the opportunity to recruit patients at the Klinikum am Europakanal Clinic for Psychiatry, Psychotherapy, and Psychosomatics and their continued support.

## References

Bach, P., Vollstädt-Klein, S., Kirsch, M., Hoffmann, S., Jorde, A., Frank, J., et al., 2016. Increased mesolimbic cue-reactivity in carriers of the mu-opioid receptor gene OPRM1 A118G polymorphism predicts drinking outcome: a functional imaging study in alcohol dependent subjects. Eur. Neuropsychopharmacol. 25, 1128–35.

Barr, C.S., Schwandt, M., Lindell, S.G., Chen, S.A., Goldman, D., Suomi, S.J., et al., 2007. Association of a functional polymorphism in the mu–opioid receptor gene with alcohol response and consumption in male rhesus macaques. Arch. Gen. Psychiatry 64, 369–76.

Bart, G., Kreek, M.J., Ott, J., LaForge, K.S., Proudnikov, D., Pollak, L., et al., 2005. Increased attributable risk related to a functional mu–opioid receptor gene polymorphism in association with alcohol dependence in central Sweden. Neuropsychopharmacology 30, 417–22.

Beery, A.K., Zucker, I., 2011. Sex bias in neuroscience and biomedical research. Neurosci. Biobehav. Rev. 35, 565–72.

Berenbaum, S.A., Byrk, K.K., Nowak, N., Quigley, C.A., Moffat, S., 2009. Fingers as a marker of prenatal androgen exposure. Endocrinology 150, 5119–24.

Bernardi, R.E., Zohsel, K., Hirth, N., Treutlein, J., Heilig, M., Laucht, M., et al., 2016. A gene–by–sex interaction for nicotine reward: evidence from humanized mice and epidemiology. Transl. Psychiatry 6, e861.

Bond, C., LaForge, K.S., Tian, M., Melia, D., Zhang, S., Borg, L., et al., 1998. Single–nucleotide polymorphism in the human mu opioid receptor gene alters β–endorphin binding and activity: possible implications for opiate addiction. Proc. Natl. Acad. Sci. USA 95, 9608–13.

Brown, E.C.Z., Steadman, C.J., Lee, T.M., Padmanabhan, V., Lehman, M.N., Coolen, L.M., 2015. Sex differences and effects of prenatal exposure to excess testosterone on ventral tegmental area dopamine neurons in adult sheep. Eur. J. Neurosci. 41, 1157–66.

Butkevich, I.P., Barr, G.A., Vershinina, E.A., 2007. Sex differences in formalin–induced pain in prenatally stressed infant rats. Eur. J. Pain. 11, 888–94.

Chen, D., Liu, L., Xiao, Y., Peng, Y., Yang, C., Wang, Z., 2012. Ethnic–specific meta–analyses of association between the OPRM1 A118G polymorphism and alcohol dependence among Asians and Caucasians. Drug Alcohol. Depend. 123, 1–6.

Esel, E., Sofuoglu, S., Aslan, S.S., Kula, M., Yabanoglu, I., Turan, M.T., 2001. Plasma levels of beta–endorphin, adrenocorticotropic hormone and cortisol during early and late alcohol withdrawal. Alcohol alcohol 36, 572–6.

Garbutt, J.C., Kranzler, H.R., O'Malley, S.S., Gastfriend, D.R., Pettinati, H.M., Silverman, B.L., et al., 2005. Efficacy and tolerability of long–acting injectable naltrexone for alcohol dependence: a randomized controlled trial. JAMA 293, 1617–25.

Ghahramani, N.M., Ngun, T.C., Chen, P.Y., Tian, Y., Krishnan, S., Muir, S. et al., 2014. The effects of perinatal testosterone exposure on the DNA methylome of the mouse brain are late–emerging. Biol. Sex Differ. 5, 8.

Gianoulakis, C., Dai, X., Brown, T., 2003. Effect of chronic alcohol consumption on the activity of the hypothalamic–pituitary–adrenal axis and pituitary β–endorphin as function of alcohol intake, age, and gender. Alcohol Clin. Exp. Res. 27, 410–423.

Goel, N., Workman, J.L., Lee, T.T., Innala, L., Viau, V., 2014. Sex differences in the HPA axis. Compr. Physiol. 4, 1121–55.

Hammarberg, A., Jayaram–Lindström, N., Beck, O., Franck, J., Reid, M.S., et al., 2009. The effects of acamprosate on alcohol–cue reactivity and alcohol priming in dependent patients: a randomized controlled trial. Psychopharmacology (Berl.) 205, 53–62.

Heilig, M., Goldman, D., Berrettini, W., O'Brien, C.P., 2011. Pharmacogenetic approaches to the treatment of alcohol addiction. Nature Reviews Neuroscience 12, 670–684.

Huber, S., Zoicas, I., Reichel, M., Mühle, C., Büttner, C., Ekici, A., et al., in press. Prenatal androgen-receptor activation determines adult alcohol and water drinking in a sex-specific way. Addict. Biol. doi: 10.1111/adb.12540.

Kleinjan, M., Rozing, M., Engels, R.C., Verhagen, M., 2015. Co–development of early adolescent alcohol use and depressive feelings: the role of the mu–opioid receptor A118G polymorphism. Development and Psychopathology 27, 915–25.

Kornhuber, J., Erhard, G., Lenz, B., Kraus, T., Sperling, W., Bayerlein, K., et al., 2011. Low digit ratio 2D:4D in alcohol dependent patients. PLoS One 6, e19332.

Kornhuber, J., Zenses, E.M., Lenz, B., Stoessel, C., Bouna-Pyrrou, P., Rehbein, F., et al., 2013. Low 2D:4D values are associated with video game addiction. PLoS One 8, e79539.

Kranzler, H.R., Gelernter, J., O'Malley, S., Hernandez-Avila, C.A., Kaufman, D., 1998. Association of alcohol or other drug dependence with alleles of the mu opioid receptor gene (OPRM1). Alcohol Clin. Exp. Res. 22, 1359–62.

Lee, K.W., Abrahamowicz, M., Leonard, G.T., Richer, L., Perron, M., Veillette, S., et al., 2015. Prenatal exposure to cigarette smoke interacts with OPRM1 to modulate dietary preference for fat. J. Psychiatry Neurosci. 40, 38–45.

Lenz, B., Mühle, C., Braun, B., Weinland, C., Bouna-Pyrrou, P., Behrens, J. et al., 2017. Prenatal and adult androgen activities in alcohol dependence. Acta. Psychiatr. Scand. 136, 96–107.

Mague, S.D., Isiegas, C., Huang, P., Liu-Chen, L.Y., Lerman, C., Blendy, J.A., 2009. Mouse model of *OPRM1* (A118G) polymorphism has sex–specific effects on drug–mediated behavior. Proc. Natl. Acad. Sci. USA 106, 10847–52.

Marchesi, C., Chiodera, P., Ampollini, P., Volpi, R., Coiro, V., 1997. Beta–endorphin adrenocorticotropic hormone and cortisol secretion in abstinent alcoholics. Psychiatry Res. 72, 187–94.

Müller, K.U., Mennigen, E., Ripke, S., 2013. Altered reward processing in adolescents with prenatal exposure to maternal cigarette smoking. JAMA Psychiatry 70, 847–56.

Muschler, M.A., Lenz, B., Hillemacher, T., Kraus, C., Kornhuber, J., Frieling, H., et al., 2014. CAGn repeat of the androgen receptor is linked to proopiomelanocortin promoter methylation–relevance for craving of male alcohol–dependent patients Psychopharmacology (Berl.) 231, 2059–66.

O'Malley, S.S., Robin, R.W., Levenson, A.L., GreyWolf, I., Chance, L.E., Hodgkinson, C.A., et al., 2008. Naltrexone alone and with sertraline for the treatment of alcohol dependence in Alaska natives and non–natives residing in rural settings: a randomized controlled trial. Alcohol Clin. Exp. Res. 32, 1271–83.

Oslin, D.W., Berrettini, W., Kranzler, H.R., Pettinati, H., Gelernter, J., Volpicelli, J.R., et al., 2003. A functional polymorphism of the mu–opioid receptor gene is associated with naltrexone response in alcohol–dependent patients. Neuropsychopharmacology 28, 1546–52.

Racz, I., Schürmann, B., Karpushova, A., Reuter, M., Cichon, S., Montag, C., et al., 2008. The opioid peptides encephalin and beta–endorphin in alcohol dependence. Biol. Psychiatry 64, 989–97.

Ramchandani, V.A., Umhau, J., Pavon, F.J., Ruiz-Velasco, V., Margas, W., Sun, H., et al., 2011. A genetic determinant of the striatal dopamine response to alcohol in men. Mol. Psychiatry 16, 809–817.

Ray, L.A., Hutchison, K.E., 2004. A polymorphism of the mu–opioid receptor gene (OPRM1) and sensitivity to the effects of alcohol in humans. Alcohol Clin. Exp. Res. 28, 1789–95.

Ray, L.A., Bujarski, S., MacKillop, J., Courtney, K.E., Monti, P.M., Miotto, K., 2013. Subjective response to alcohol among alcohol dependent individuals: effects of the mu–opioid receptor (OPRM1) gene and alcoholism severity. Alcohol Clin. Exp. Res. 37, E116–124.

Schacht, J.P., Randall, P.K., Latham, P.K., Voronin, K.E., Book, S.W., Myrick, H., et al., 2017. Predictors of naltrexone response in a randomized trial: reward–related brain activation, *OPRM1* genotype, and smoking status. Neuropsychopharmacology, [Epub ahead of print].

Schwantes-An, T.H., Zhang, J., Chen, L.S., Hartz, S.M., Culverhouse, R.C., Chen, X., et al., 2016. Association of the OPRM1 variant rs1799971 (A118G) with non–specific liability to substance dependence in a collaborative *de novo* meta–analysis of European–Ancestry cohorts. Behav. Genet. 46, 151–69.

Stuppaeck, C.H., Barnas, C., Falk, M., Guenther, V., Hummer, M., Oberbauer, H. et al., 1994. Assessment of the alcohol withdrawal syndrome – validity and reliability of the translated and modified Clinical Institute Withdrawal Assessment for Alcohol scale (CIWA–A). Addiction 89, 1287–92.

van den Wildenberg, E., Wiers, R.W., Dessers, J., Janssen, R.G., Lambrichs, E.H., Smeets, H.J., et al., 2007. A functional polymorphism of the mu–opioid receptor gene (OPRM1) influences cue–induced craving for alcohol in male heavy drinkers. Alcohol Clin. Exp. Res. 31, 1–10.

Veening, J.G., Gerrits, P.O., Barendregt, H.P., 2012. Volume transmission of beta–endorphin via the cerebrospinal fluid; a review. Fluids Barriers CNS 9, 16.

Zalewska-Kaszubska, J., Czarnecka, E., 2005. Deficit in beta–endorphin peptide and tendency to alcohol abuse. Peptides 26, 701–5.

Zhang, Y., Wang, D., Johnson, A.D., Papp, A.C., Sadée, W., 2005. Allelic expression imbalance of human mu opioid receptor (OPRM1) caused by variant A118G. J. Biol. Chem. 280, 32618–24.

Zheng, Z., Cohn, M.J., 2011. Developmental basis of sexually dimorphic digit ratios. Proc. Natl. Acad. Sci. USA 108, 16289–94.

